# Shift and adapt: the costs and benefits of karyotype variations

**DOI:** 10.1101/025171

**Authors:** Aleeza C. Gerstein, Judith Berman

**Author notes:** Corresponding author: Judith Berman.

## Abstract

Variation is the spice of life or, in the case of evolution, variation is the necessary material on which selection can act to enable adaptation. Karyotypic variation in ploidy (the number of homologous chromosome sets) and aneuploidy (imbalance in the number of chromosomes) are fundamentally different than other types of genomic variants. Karyotypic variation emerges through different molecular mechanisms than other mutational events, and unlike mutations that alter the genome at the base pair level, rapid reversion to the wild type chromosome number is often possible. Although karyotypic variation has long been noted and discussed by biologists, interest in the importance of karyotypic variants in evolutionary processes has spiked in recent years, and much remains to be discovered about how karyotypic variants are produced and subsequently selected.

## Introduction

Fungal microbes have emerged as important taxa to study the effect of karyotypic variation on phenotype and fitness (reviewed in [1]). Ploidy variation exists among, as well as within, fungal species and ploidy shifts continue to emerge in surprising places. Variation in ploidy among nuclei that share a common cytoplasm was recently reported in *Ashbya gossypii*, an otherwise haploid organism [2**]. *Candida albicans*, long considered an obligate diploid, recently was found to generate haploids following passage through a mouse model, and following exposure to the antifungal drug fluconazole [3,4]. Similarly, rare tetraploids have been sampled from clinical isolates of *C. albicans* [5,6], and polyploid “titan” cells were recently identified in the typically-haploid basidiomycete *Cryptococcus neoformans* [7,8] (reviewed in [9]). Aneuploid variants also have been documented at appreciable frequencies in many fungal taxa in a diversity of circumstances: within the *Saccharomyces. cerevisiae* deletion collection constructed by transformation [10], following drug stress in *C. albicans* [11] and *C. neoformans* [12], in meiotic spores of *Cryptococcus lusitaniae* [13] and through unisexual mating in *C. neoformans* [14]. Furthermore, it remains possible that the true proportion of karyotypic variants is even higher than currently appreciated, as variants are rapidly eliminated *ex vivo* due to the growth conditions typically used to propagate cultures in the laboratory.

Here we focus on karyotypic variants that arise mitotically, within a single organism, rather than those formed through conventional means involving sexual conjugation followed by meiotic reductive divisions. We first discuss variations in whole genome ploidy level and then changes due to aneuploidy, focusing on the mechanisms that produced the variation and then on the degree to which they provide a selective advantage under different growth conditions.

## Producing ploidy shifts

The mitotic mechanisms by which ploidy variants are produced are dependent on the direction of change. Increases occur through endoreplication, a process in which the genome replicates, but the cell containing it fails to complete mitosis and cytokinesis. Decreases in ploidy occur via a cryptic chromosome loss mechanism, often, if not always, through aneuploid intermediates [3,15,16]. There remains no known mitotic program to undertake a full euploid chromosome set reduction (‘concerted chromosome loss’ [17]), yet the rate at which some reductions in ploidy have been documented in the laboratory indicates that it can not be attributed solely to selection acting on individual chromosome loss events. Ploidy transitions frequently occur within relatively short timescales; they are often detected following fitness assay tests (24-72 h) and short-term experimental evolution passaging (< 200 generations). Convergence towards the historical ploidy level has now been documented in *S. cerevisiae* [15,18,19], *C. albicans* [3,16], *C. tropicalis* [20], and *C. neoformans* (Gerstein, Fu et al., in review). The force behind this ‘ploidy-drive’ remains cryptic, as there have been few identified fitness differences between ploidy-variant individuals sampled from the same population [21]. The presence of ploidy variants and the rate of ploidy transition is strongly influenced by the environment [2,15,17], yet the link(s) between genotype, phenotype and fitness remain murky.

## Does ploidy state provide a selective advantage?

An ongoing topic of debate in evolutionary biology is why one ploidy level should be beneficial over another. As all genomes are subjected to the same basic processes, why is there not a singular “best” ploidy level employed across the tree of life? This question has been addressed using fungal microbes that can be manipulated to generate isogenic ploidy series, thereby distinguishing ploidy from differences in genetic background. Predictions generally fall into two categories: short-term influences of ploidy on ecological tolerance and survival, mediated through physiological differences; and long-term (evolutionary) influence of ploidy on the rate of adaptation, mediated through the genetic basis of beneficial mutations (discussed in detail in [22]).

Physiologically, ploidy is frequently correlated with differences in the cell surface area:volume (SA:V) ratio, which decreases as ploidy and cell size increase. Lower ploidy (and a higher SA:V ratio) is predicted to be beneficial under nutrient limitation, yet detrimental when toxins are present in the environment. Overall, these predictions received only mixed support from empirical tests, as highlighted by Zorgo *et al.* [23**], who compared many haploid and diploid strains of *S. cerevisiae* and S. *paradoxus*. They found that although ploidy × environment interactions were frequent and often evolutionarily conserved, there was not an overall growth advantage for diploids in toxins, or for haploids under nutrient limitation or environmental stress. Thus, no single catch-all explanation accounts for why one ploidy level is ecologically beneficial over another. This result may not be satisfying, but it likely reflects the reality of biology: a large number of factors usually influence growth and survival.

Recent technological advances, in particular flow cytometry coupled with next generation sequencing, now enable researchers to track the genetic basis of adaptation and to test the influence of ploidy on the mutations selected during adaptation. When beneficial mutations arise in diploids or polyploid individuals, their effects are often at least partially masked by the wild-type allele(s) that remain. Thus, haploids may be able to acquire a greater diversity of mutations (as in [24]) or to access different types of mutations. This was recently shown for lines adapting to nutrient limitation: beneficial loss-of-function mutations, which are often recessive, arose more frequently in haploid lines, while diploid lines were enriched for mutations that increase gene expression (and are likely at least partially expressed in heterozygous form, [25]). Thus haploids, which fully express all beneficial mutations, might be expected to adapt faster than diploids. However, depending on the environmental challenge, this may [26], or may not [19], be the case. An increase in ploidy also increases the mutation rate [19], and, if selection is strong, loss of heterozygosity can rapidly convert heterozygous mutations to a homozygous state [27], thereby reducing the difference in the rate of adaptation between haploids (where the effect of recessive beneficial mutations are immediately felt) and higher ploidy levels. Finally, mutations may also have different effect sizes in different ploidy backgrounds, even when present in homozygous form [28]. Taken together, it appears that the relationship between ploidy and adaptation depends heavily on the environment and the particulars of the available mutations. Consistent with this idea, a recent genome-wide tour-de-force by Dunham and co-workers distinguished the effects of aneuploidy from those of mutations acquired in aneuploid strains using a clever system to generate extra copies of chromosome arm segments. They found that aneuploidies could provide adaptive advantages under specific environmental conditions and that the advantages were clearly related to the genes included on the aneuploid segments [29**].

## Slipping into (and out of) aneuploidy

Aneuploidy arises through nondisjunction of single, or a few, chromosomes, and provides a fundamentally different class of adaptive genomic change than point mutations (Figure 1). As chromosome nondisjunction occurs several orders of magnitude more frequently than point mutations, and aneuploidy is more prevalent than shorter range copy number variations, small insertions, or small deletions [30], aneuploidy provides a rapid mechanism to generate diversity upon which selection can act (Figure 1b). Aneuploidy may also act as a stepping-stone to other types of genic mutations if the environmental stress is maintained for many generations. Similar to changes in ploidy, aneuploidy may increase the rate that beneficial mutations can arise within the genome (Figure 1c), either on the chromosome that is aneuploid, or through a general increase in genomic instability. Once a chromosome imbalance is present, the fidelity of chromosome segregation appears to be reduced. If additional beneficial point mutations are then acquired, selection to maintain the aneuploidy will be lost. When the environmental stress is removed, aneuploidy also can be lost more rapidly than other genomic mutations, via a chromosome non-disjunction event (Figure 1d). Thus, aneuploidy provides a highly flexible mechanism to rapidly adapt to a changing environment.

**Figure 1.**
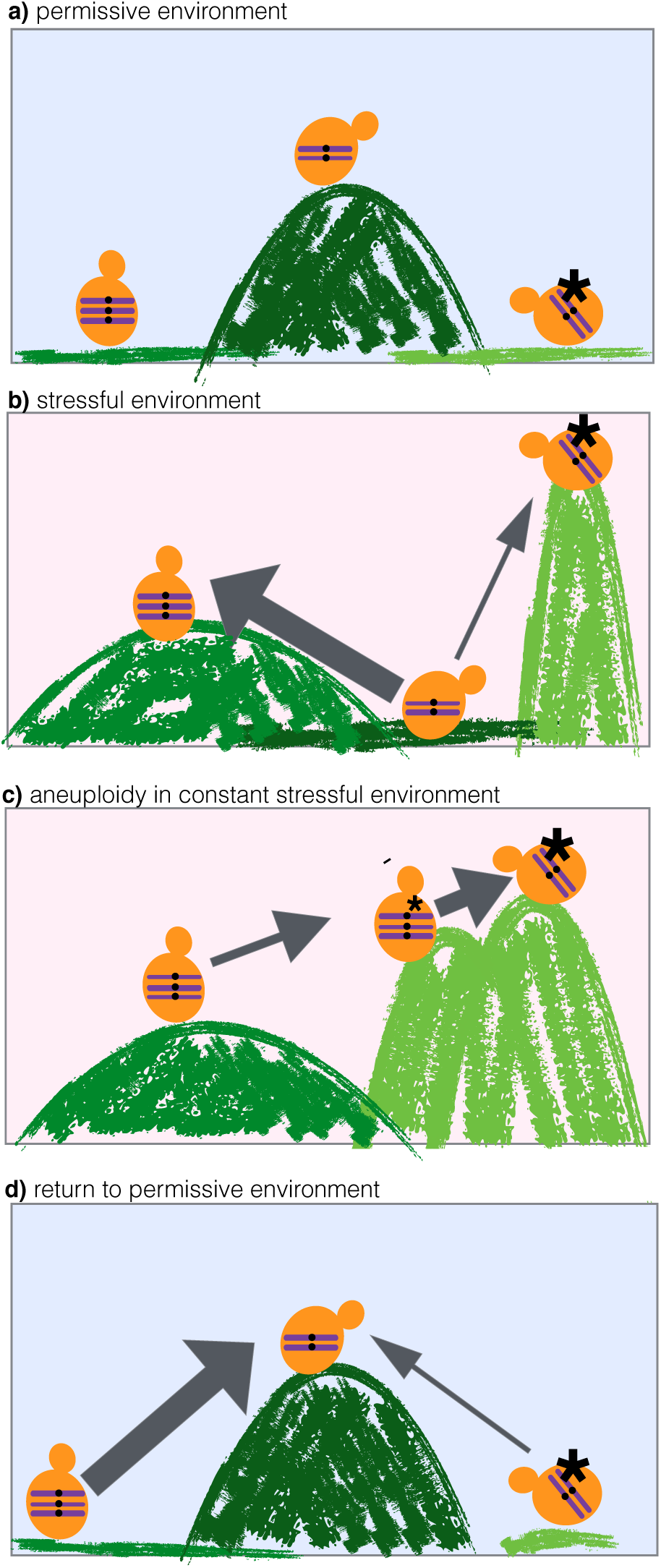
Aneuploidy can be rapidly selected for or against in permissive environments (blue background) and stressful environments (pink background). (a) In a permissive environment the wild-type genotype is more fit (occupies a higher peak on the landscape) than either an aneuploid strain or a mutant strain. (b) When in a stress environment, the wild-type genotype is no longer the most fit. Multiple types of mutations can confer a fitness advantage, including a particular aneuploidy and a genic mutation. The width of the arrows indicates the mutational frequency; aneuploidy arises more frequently than any particular genic mutation. (c) If the stress environment is maintained over time, aneuploidy can act as a bridge to genic mutations. The mutation rate is increased in aneuploids, due to the extra chromosome copy, and the aneuploidy can be easily lost once a beneficial mutation arises. (d) If the environment returns to being permissive, reversion of an aneuploid chromosome via chromosome gain or loss can occur with high frequency. Reversion of genic mutations is a much rarer event.

Chromosomes in aneuploid cells are more prone to nondisjunction events than are chromosomes in euploid cells [16,31,32]. Thus, once the initial aneuploidy is present the rate with which additional aneuploidy events appear is increased. The rate of aneuploid formation also increases during stress exposure. Within the first 12-24 hours of drug exposure, over 20% of *C. albicans* cells undergo unusual mitotic divisions that yield tetraploids/heterokaryons that then frequently give rise to aneuploids and, rarely, haploids [33**], [4]. Hsp90 may also play an important role in aneuploid formation under stress conditions, as higher chromosome non-disjunction rates have been observed under stressors that demand increased levels of Hsp90 and that compromise kinetochore assembly [34]. Thus, extreme stress can lead to unconventional mitotic mechanisms of ploidy change under strong selective pressure that inhibits the growth of the original parent [33**]. Taken together, the appearance of aneuploids appears to be a natural consequence of cell cycle progression defects caused by stress itself, reminiscent of fitness associated recombination [35], stress-induced mutagenesis in bacteria (reviewed in [36]) and stress-induced loss of heterozygosity in yeasts [37].

## Aneuploidy and stress: is the benefit worth the cost?

Aneuploidy has the potential to affect phenotypic variation on both a general and specific level. A general signature of aneuploidy that is independent of the specific chromosome(s) involved has been proposed to involve slower progression through G1-phase [38,39] and elevated levels of proteotoxic stress relative to euploids [40]. Many aneuploid *S. cerevisiae* strains exhibit this feature, albeit to different degrees, but some do not [38]. Alterations in mRNA and protein levels generally correspond to the alterations in DNA copy number [41-44],though this effect may be buffered for certain genes and/or strain backgrounds [45] or be mitigated by post-translational regulatory effects on unlinked genes and their products [46]. Specific phenotypes linked to extra gene copies are often evident, though the effect of aneuploidy may be realized by the combined influence of multiple copy number changes [29**,47]. While the majority of possible aneuploidies likely have deleterious phenotypes (reviewed in [48]), selection should efficiently eliminate these deleterious aneuploidies from large populations, thereby enabling beneficial aneuploidies to remain.

If a favourable genotype can be acquired through aneuploidy, this may be expected to be a frequent route to adaptation, due to the higher rate of aneuploid formation relative to other types of mutations. Indeed, a high rate of beneficial aneuploidy has been identified under glucose limitation [49], drug stress [50], copper adaptation [51], adaptation to sulphate limitation [18,29**] and as a solution for *gal7Δ* strains that require a beneficial mutation to grow in galactose [52]. Furthermore, aneuploidy appears to be a more common event in diploids than in haploids [18], perhaps because the deleterious effects of an extra gene copy are less dramatic for diploids than for haploids. The propensity for specific aneuploidies to be selected as a result of specific environmental pressures has been exploited to measure chromosome gain rates in different genetic backgrounds [53]. The reliable production of aneuploids can also be exploited to set an “evolutionary trap”, by use of a drug that first enriches the population for a particular aneuploidy and a second drug that specifically inhibits the growth of cells that carry this aneuploidy.[54**]. As a general principle, aneuploidies are likely to incur a cost in non-selective environments, with the magnitude of this cost (and thus the stability of the aneuploidy) dependent on interaction between the environment, genes that are present in the aneuploidy, and the genetic background [44,51,55,56].

The degree of stress, independent of the environment, may also play a role in selecting for or against aneuplodies. Indeed, the degree to which fitness varied among otherwise isogenic aneuploid strains of *S. cerevisiae* scaled with the degree of stress across a range of conditions, suggesting that there may be a functional relationship between the degree of stress applied and the degree of phenotypic variation (including beneficial variation) [54]. This has important implications for therapeutic strategies, as it suggests that more severe drug treatments may stimulate more population diversity and may lead to progeny with a higher potential to adapt to the drug. The reverse may also be true, however, as aneuploidy may be seen less frequently under stressful conditions in multinucleate filamentous fungi [2**].

As a result of both the general cost of aneuploidy and the specific chromosome × environment trade-offs, aneuploidy is likely to be a temporary evolutionary solution, until more stable and specific genetic solutions arise. For example, *S. cerevisiae* cells that evolved under mild temperature stress for 450 generations first acquired an extra copy of Chr3, which facilitated faster growth at elevated temperature [57**]. Later, following further passage under elevated temperature, the aneuploidy was lost. A phenomenon reminiscent of this appeared in series of *C. albicans* clinical isolates from patients, where Chr5 aneuploidy often preceded the emergence of highly drug-resistant variants [1,58]. As aneuploidy is easily reversible, it may also function as a bet-hedging mechanism for toggling between two phenotypic states, as was seen for colony morphology phenotypes of *S. cerevisiae* isolates [2**,59*].

## Conclusion

We propose that karyotypic variation can be both a non-selected cause and an adaptive response to stress that enables phenotypic variation to appear rapidly within populations. This is demonstrated by the response of *C. albicans* in the presence of the antifungal drug fluconazole: aneuploidy arises as a consequence of exposure to a drug, which stimulates the formation of multinucleate/tetraploid cells that produce aneuploid cells at a high rate [3*,4,33**]. Specific forms of aneuploidy are also beneficial under fluconazole stress, and thus are maintained as a result of selection [12,60]. Importantly, karyotypic variants tend to be easily reversible, and thus under unstable environmental conditions they can rapidly appear and disappear from populations.

Karyotypic variants do not always incur a high fitness cost and relatively stable variants have been found during short and long term laboratory experimental evolution experiments, in a number of series of isolates from patients, and in nature [18,29**,44,61-63]. However, due to their transient nature, sampling results and clinical sampling practices may under-estimate the true number of karyotypic variants. Similarly, in laboratory evolution experiments we likely miss a large degree of variation by focusing efforts primarily on the last, usually arbitrary, time-point. Finally, it is important to note that the majority of evolution experiments with *S. cerevisiae* have used haploid isolates that can undergo only chromosome gains, but not chromosome losses, and that may be less tolerant of additional chromosome copies than diploids. Not only is the appearance of aneuploidy likely to be more frequent in diploids or polyploids than in haploids [18], but aneuploidy well may have different fitness consequences as well [19*].

In conclusion, ploidy and aneuploidy variants have the potential to arise frequently within populations, and may facilitate rapid adaptation. The emerging view is that the influence of aneuploidy is context dependent, and the fitness consequences depend on all aspects of a given circumstance: the environment, the genetic background, and the genes that are present in aneuploidy.

We apologize to the authors of many important studies whose work we did not have the space to discuss due to the concise nature of this article format.

## Acknowledgements

We thank Anna Selmecki for helpful discussions. This work was supported by the People Programme (Marie Curie Actions) the European Union’s Seventh Framework Programme (FP7/2007-2013) REA grant agreement number 303635; by an ERC Advanced Award, number 340087, RAPLODAPT, grants from the Israel Science foundation (340/13.), and by the National Institute of Allergy and Infectious Disease (R01AI075096 and R01AI0624273) to JB. ACG was supported by fellowship awards from the National Sciences and Engineering Research Council of Canada, the Canadian Institutes for Health Research, and the Azrieli Foundation.

